# Streptococcus pyogenes Cas9 displays biased one-dimensional diffusion on dsDNA to search for a target

**DOI:** 10.1101/754564

**Authors:** Mengyi Yang, Lijingyao Zhang, Ruirui Sun, Weixiong Zhong, Yuzhuo Yang, Chunlai Chen

## Abstract

The RNA-guided Streptococcus pyogenes Cas9 (spCas9) is a sequence-specific DNA endonuclease that works as one of the most powerful genetic editing tools. However, how the Cas9 locates its target among huge amounts of dsDNAs remains controversial. Here, combining biochemical and single-molecule assays, we revealed that Cas9 uses both three-dimensional and one-dimensional diffusion to find its target. We further observed a surprising biased one-dimensional diffusion of Cas9 from 3’ to 5’ end of the non-target strand under physiological salt condition, whereas low ionic concentration or mutations on PAM recognition residues induce unbiased one-dimensional diffusion of Cas9 along dsDNA. We quantified the diffusion length of 27 bp, which accelerates the target search efficiency of Cas9 by ∼ 10 folds. Our results reveal a unique searching mechanism of Cas9 at physiological salt conditions, and provide important guidance for both in-vitro and in-vivo applications of Cas9.

## Introduction

CRISPR (Clustered Regularly Interspaced Short Palindromic Repeats) and CRIPSR-associated (Cas) protein function as adaptive immune system to protect the bacteria and archaea from invading bacteriophages and plasmids (Barrangou et al., 2007; Brouns et al., 2008; Marraffini, 2015; Wiedenheft et al., 2012; Wright et al., 2016). In the class two type II system, the signature Cas9 endonuclease is guided by a chimeric single-guide RNA (sgRNA) to find, bind and cleave a 20-bp-length complementary DNA flanked by a PAM (protospacer adjacent motif) site (Anders et al., 2014; Jinek et al., 2012; Jinek et al., 2014; Singh et al., 2016; Szczelkun et al., 2014). CRISPR-Cas9 has been exploited as one of the most powerful tools in genomic engineering in living cells. (Barrangou and Doudna, 2016; Cong et al., 2013; Doudna and Charpentier, 2014; Hsu et al., 2014; Knott and Doudna, 2018).

At the cellular level, Cas9 has to search among huge amounts of dsDNA to find its target with high accuracy and efficiency. After initial interaction with dsDNA, Cas9/sgRNA complex unstably bind at non-PAM site and is quickly released, whereas PAM interactions at cognate target trigger local melting of dsDNA at the PAM-adjacent nucleation site to promote further R-loop formation and extension (Anders et al., 2014; Sternberg et al., 2014; Szczelkun et al., 2014). Stable Cas9/sgRNA/dsDNA ternary complex forms when PAM-proximal 8 ∼ 12 nt region (seed) is fully matched towards the guide RNA. Various efforts have previously been employed to understand how Cas9 searches for its target on dsDNA. DNA curtain experiments proposed a random collision (three-dimensional diffusion, 3D diffusion) model of Cas9 to search for its targets (Sternberg et al., 2014). With improved spatial resolution, single-molecule fluorescence resonance energy transfer (smFRET) assays under low salt conditions capture one-dimensional (1D) diffusion of Cas9 among multiple PAM sites on the same dsDNA (Globyte et al., 2019). However, how Cas9 finds its target under physiological salt conditions remains largely unknown.

Here, by combining ensemble biochemical and single-molecule fluorescent assays, we discovered that the Streptococcus pyogenes Cas9 (SpCas9) combined both 3D and 1D diffusion to find its targets. Surprisingly, Cas9 displayed biased 1D diffusion from 3’ to 5’ of non-target strand to search for PAM sites under physiological salt condition. Low salt concentration or mutations on PAM recognition residues switched Cas9 to unbiased 1D diffusion. We further quantified that the 1D diffusion length of Cas9 on dsDNA is ∼ 27 bp, which increased the rate of target search by 10 folds or more. Therefore, to ensure rapid target search, the accessibility of target sites and their nearby Cas9 binding sites are both required. Our discoveries provided new guidance to design and optimize Cas9 target sites for in-vitro and in-vivo applications.

## Results

### One-dimensional (1D) diffusion of Cas9 on dsDNA

To investigate target search mechanisms of spCas9 under close to physiological condition, 150 mM KCl was used, unless stated otherwise. Time-dependent in-vitro cleavage assay was performed towards a series of dsDNAs which were linearized from the same plasmid containing a single cleavage site (Fig. 1a, Table S1). The target site was centered in a linearized dsDNA fragment, whose length varied from 30 bp to 2188 bp. When needed, a linearized non-target fragment was supplied to maintain identical total length and sequence of dsDNAs. Cleavage rates of Cas9/sgRNA towards target-containing dsDNAs of different length were resolved and quantified (Fig. 1b-c, Table S2). If Cas9 locates the target simply by 3D diffusion, the cleavage rates towards target-containing dsDNAs should be independent of DNA length. Clearly, apparent cleavage rates increased significantly when the length of dsDNA increased from 30 bp to 120 bp and remained almost constant when target-containing dsDNA were 120 bp or longer. Therefore, we speculate that extended dsDNA fragments flanking target site accelerate target searching process via a previously proposed facilitated diffusion model (Berg et al., 1981; Gerland et al., 2002; Halford and Marko, 2004; von Hippel and Berg, 1989), which involves initial associations at random DNA sites, followed by 1D diffusion of Cas9 to locate its specific targeting sites.

**Figure 1.**
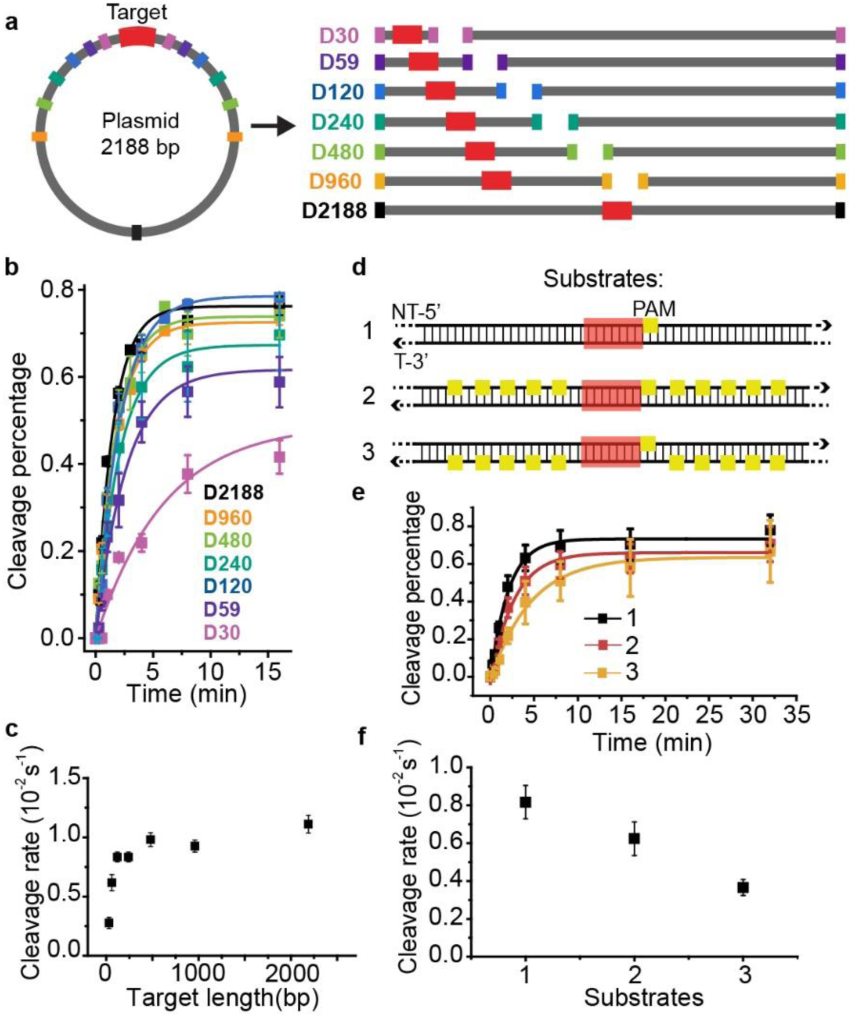
Time-dependent in-vitro cleavage asssay revealed 1-D diffusion of Cas9 during target search. **a)** Experimental design of linearized dsDNAs, which were generated from a 2188-bp-length plasmid containing a single target site. The target site (shown in red) was centered in linearized dsDNA fragments, who were named after their length from D30 to D2188. When needed, linearized supplemental fragments containing no target site were supplied to maintain identical total length and sequence. **b)** Time-dependent in-vitro cleavage assay of Cas9/sgRNA towards the sequences designed in **a**, resolved by agarose gel or denaturing polyacrylamide electrophoresis. Percentage of cleaved DNA was visualized by GE typhoon FLA9500 imaging system and quantified by exponential fit. **c)** Cleavage rates extracted from curves shown in **b. d)** Three dsDNAs of 617 bp in length containing a centered target site (target sesquence is colored red with a PAM in yellow) without other PAM sites (substrate 1), with 10 PAMs flanked on non-target strand (substrate 2), and with 10 PAMs flanked on target strand (substrate 3). **e)** Time-dependent in-vitro cleavage assay of Cas9/sgRNA towards DNA constructs in **d. f)** Cleavage rates extracted from curves shown in **e**. Errors were S.E.M. from three or more replicates.

Cas9 is capable of transient binding to PAM sites in the absence of heteroduplex formation (Mekler et al., 2017). In agreement with previous reports (Anders et al., 2014; Mekler et al., 2017; Sternberg et al., 2014), multiple PAM sites surrounding the cleavage site were able to significantly inhibit apparent cleavage rate (Figs. 1d-f, Tables S1-S2). Notably, target flanked by ten PAM sites on the non-target (NT) strand was cleaved faster than that flanked by ten PAMs on the target (T) strand (Figs. 1d-f). Such phenomena argued against the model that Cas9 uses only 3D diffusion to search for its target. We proposed that, when Cas9 diffuses one-dimensionally on dsDNA, it searches for PAM sites and recognizes target sites on one strand. To search on the other strand, Cas9 has to dissociate and bind dsDNA in the reverse orientation. Consistent with this model, only Cas9 interacting with PAM sites on NT strand can diffuse one-dimensionally to the nearby cleavage site without dissociation, which caused faster cleavage rate of the target flanked by NT PAM sites.

### Biased 1D diffusion under physiological buffer condition

Next, we placed target sites at different locations of 59-bp-length dsDNAs. Surprisingly, target site located close to 5’ end of NT strand (D8-PAM-28) was cleaved 5 folds faster than that located close to 3’ end of NT strand (D28-PAM-8, Fig. 2, Tables S1-S2). It suggested that Cas9 displays biased 1-D diffusion from 3’ end of NT strand to 5’ end. Similar behaviors were captured when a different set of dsDNAs were used (Tables S1-S2), supporting that the biased 1D diffusion of Cas9 is DNA sequence independent.

**Figure 2.**
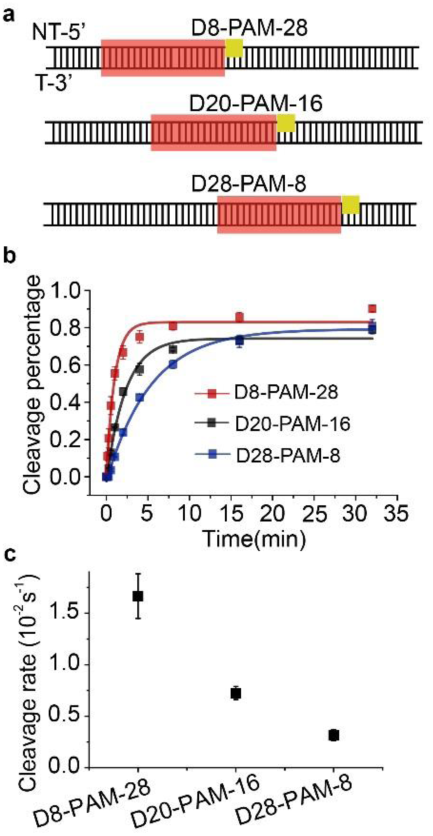
Time-dependent in-vitro cleavage asssay revealed biased 1-D diffusion of Cas9. **a)** Design of three 59-bp dsDNAs, whose target sequences (shown in red) and PAMs (shown in yellow) were placed at different locations. dsDNAs were named after the location of target and PAM sites relative to the 5’ and 3’ ends of the NT strand, respectively. For example, dsDNA D28-PAM-8 indicates that there are 28 base pairs between target site and 5’ end and 8 base pairs between PAM site and 3’ end. The following dsDNAs were named after the same nomenclature. **b)** Time-dependent in-vitro cleavage assay of Cas9/sgRNA towards dsDNAs shown in **a**. Percentage of cleaved DNA was resolved by denaturing polyacrylamide electrophoresis, visualized by GE typhoon FLA9500 imaging system and quantified by single exponential fits. **c)** Cleavage rates extracted from curves shown in **b**. Errors were S.E.M.

### Quantification of biased 1D diffusion under physiological buffer condition

Single-molecule fluorescence assays were used to quantify biased 1D diffusion of Cas9. Fluorophore labeled Cas9/sgRNA complexes were flowed into reaction chamber containing immobilized biotinylated dsDNA (Figs. 3a, c and e, Table S3). Time from injection of Cas9 to the appearance of individual stable binding events were recorded, whose distributions were fitted by single exponential decay. Their reaction time constants and appearance rates were subsequently determined (Figs. 3b, d and f).

**Figure 3.**
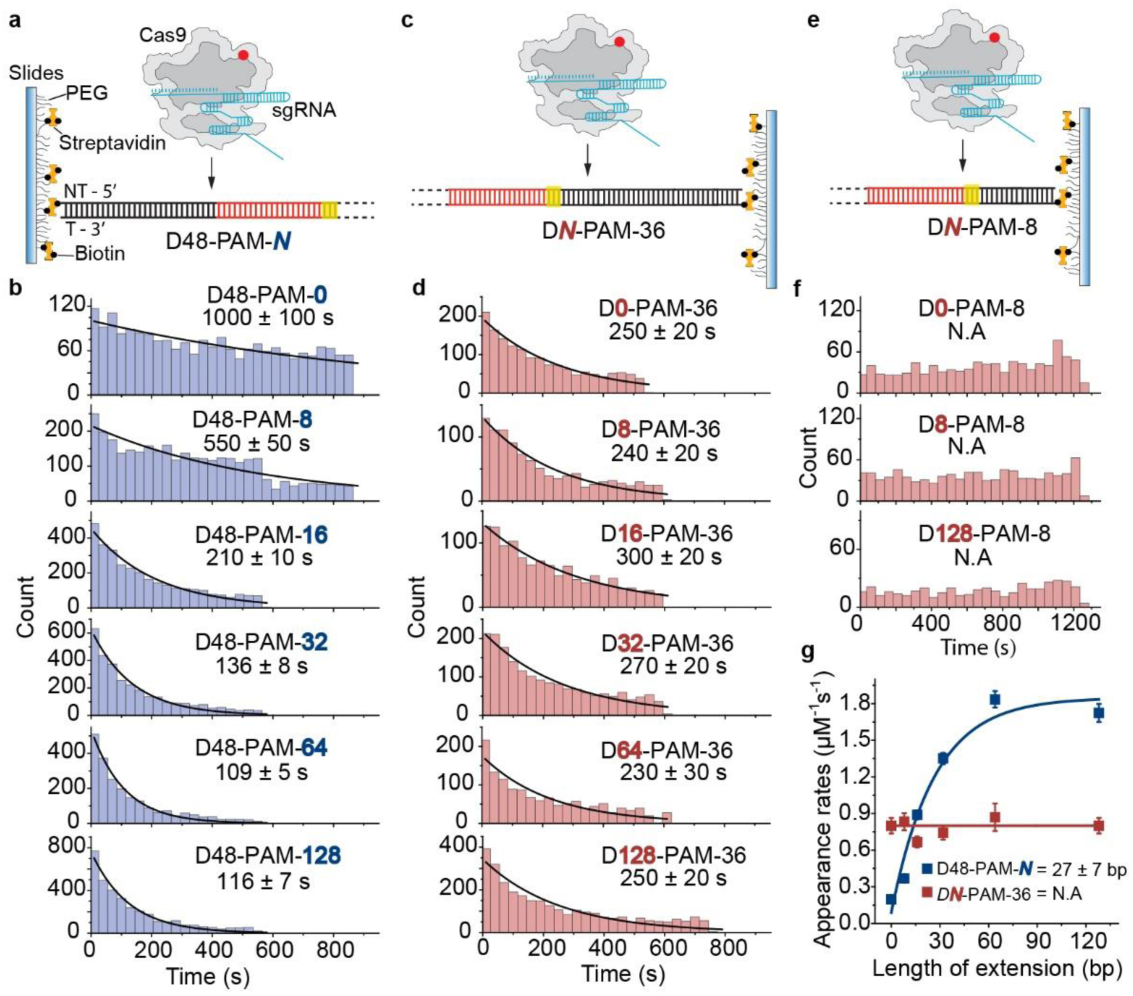
Single-molecule fluorescence assay quantifying biased 1D diffusion of Cas9. **a)** Scheme of single-molecule fluorescence assay of capturing apparent target search rates of Cas9 on D48-PAM-*N* DNAs. Biontinylated dsDNAs were immobilized on PEG passivated microscope glass slide. Time from injecting pre-incubated labeled Cas9/sgRNA onto immobilized dsDNAs until stable binding events occurred were recorded. **b)** Distributions of appearance time of Cas9 on D48-PAM-*N* DNAs, which were fitted by single exponential decay. **c)** Scheme of capturing apparent target search rates of Cas9 on D*N*-PAM-36 DNAs. **d)** Distributions of appearance time of Cas9 on D*N*-PAM-36 DNAs. **e)** Scheme of assays with D*N*-PAM-8 DNAs. **f)** Distributions of appearance time with D*N*-PAM-8 DNAs. **g)** Apparent target search rates of Cas9 on D48-PAM-*N* and D*N*-PAM-36 DNAs. Extension of dsDNA at 5’ end of NT strand had almost no effect on search rates (red), whereas extension at 3’ end of NT strand increased search rates significantly (blue). 1D diffusion length of Cas9 was estimated as 27 ± 7 bp.

For the targeting dsDNAs of the D48-PAM-*N* series, the target sites were positioned where the distance to 5’ end of NT strand remained the same while the distance to 3’ end of NT strand extended from 0 bp to 128 bp (Figs. 3a and b, Table S3). We discovered that the extended dsDNA at 3’ end of NT strand greatly facilitated target search of Cas9 by ∼ 10 folds (Fig. 3b and Table S4). On the other hand, for the D*N*-PAM-36 series (Table S3), when the distance between the target sites and 5’ end of NT strand extended from 0 bp to 128 bp, appearance rates of Cas9 on dsDNA stayed almost the same (Fig. 3d and Table S4). Similar behaviors were captured when three different sets of dsDNAs were used (Fig. S1, Tables S3-S4), which eliminated the possibility of sequence-dependent mechanisms. As a further support, when we used the D*N*-PAM-8 series, the extension of 5’ end of NT strand from 0 to 128 bp also has no effect on increasing the appearance rates of Cas9, suggesting the slow appearance rate of Cas9 to D128-PAM-8 is mainly caused by the short 8-bp extension at 3’ end of NT strand (Fig. 3f, Table S4). Together, these single-molecule assays further supported the model that Cas9 searches its targets via biased 1D diffusion from 3’ end of NT strand to 5’ end. We quantified that the bias 1D diffusion length of Cas9 was 27 ± 7 bp (Fig. 3g), which is close to the previously reported ∼ 20 bp of the diffusion length (Globyte et al., 2019).

By combining 3D and 1D diffusion, the maximal effective target search rate of Cas9 is *k*_eff_ = 1.8 ± 0.1 μM^-1^ s^-1^ under our experimental conditions (Table S4), which is close to a previously reported value (∼ 4 μM^-1^ s^-1^) (Singh et al., 2016). Furthermore, we estimated the 3D binding rate between Cas9 and one binding site on dsDNA via *k*_3D_ = *k*_eff_/*L*, in which *L* = 27 ± 7 is 1D diffusion length of Cas9 on dsDNA. Together, we have *k*_3D_ = 0.07 ± 0.02 μM^-1^ s^-1^.

### Unbiased 1D diffusion in low-salt buffer

Ionic strength is known to affect the kinetics and interactions between DNA and DNA-binding proteins (Berg et al., 1981; Bonnet et al., 2008; Lohman, 1986; Tempestini et al., 2018). To evaluate how ionic strength affects 1D diffusion of Cas9, we performed single-molecule assays shown in Fig. 3 under low-salt buffer containing 10 mM NaCl (Fig. S2). Strikingly, extension of dsDNA on either 5’ or 3’ end of NT strand increased rates of Cas9 target search, suggesting an unbiased 1-D diffusion mechanisms at low salt condition. Our findings agreed with previous smFRET measurements in the same low-salt buffer (Globyte et al., 2019).

### 1D diffusion of Cas9 variants recognizing alternative PAMs

Both our ensemble and single-molecule assays indicated that Cas9 diffuses unidirectionally on dsDNA to search and recognize PAM sites, which permits sequential Cas9/sgRNA/dsDNA complex formation and cleavage (Anders et al., 2014; Sternberg et al., 2014; Szczelkun et al., 2014). By modifying PAM-interacting residues, Cas9-VQR (D1135V/R1335Q/T1337R) and Cas9-EQR (D1135E/R1335Q/T1337R) have been engineered to recognize alternative 5’-TGAN-3’ PAM sequences (Anders et al., 2016; Hirano et al., 2016). Their 1D diffusion behaviors were examined by our single-molecule assays. Cas9-VQR displays similar biased 1D diffusion as the wild-type Cas9 (Table S5). For Cas9-EQR variant, the extended dsDNA on both 5’ and 3’ end of NT strand accelerates the appearance rates of Cas9 towards its targets (Fig. S3, Table S5), exhibiting unbiased 1D diffusion in target search process. Such phenomena suggest Cas9 residues interacting with dsDNA to recognize PAM sites affects the 1D diffusion of Cas9.

## Discussion

Our results support a model in which Cas9 combines 3D and 1D diffusion to find its target with high efficiency and accuracy (Fig. 4). Freely-diffusing Cas9/sgRNA complex interact with dsDNA through 3D random collision. Once upon interacting with dsDNA, Cas9 diffuses laterally along dsDNA (1D) for ∼ 27 bp before dissociation, unless a target site is encountered. Interestingly, Cas9 displays biased 1D diffusion from 3’ to 5’ of non-target strand to search for PAM sites under physiological salt condition. When Cas9 encounters the PAM site of its cognate target, it initiates RNA strand invasion and R-loop expansion (Gorski et al., 2017; Jiang and Doudna, 2017). On the other hand, Cas9 continues its 1D diffusion on dsDNA after transient binding with PAM sites flanking non-target sequence, which is consistent with previous reports (Globyte et al., 2019). Lastly, we quantified that 1D diffusion facilitates target search of Cas9 by ∼ 10 folds (Fig. 3 and Table S4).

**Figure 4.**
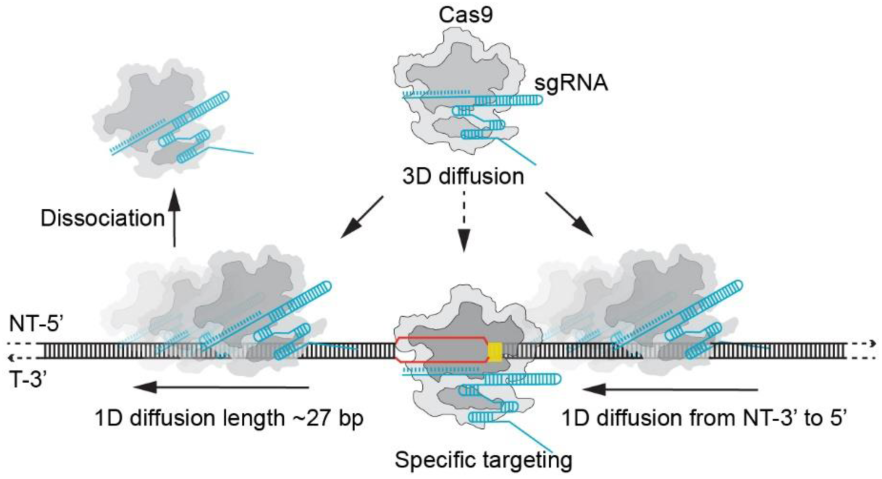
A proposed schematic model of target search mechanism by Cas9. Cas9 combines both 3D and biased 1D diffusion to find its target under physiological salt condition. Freely-diffusing Cas9/sgRNA molecule interacts with dsDNA through 3D random collision. Once upon interacting with dsDNA, Cas9 diffuses laterally along dsDNA (1D) for ∼ 27 bp before dissociation, unless a target site in encountered.

With improved time resolution of 2.5 ms per frame, we captured that the total dwell time of Cas9 on a PAM-free dsDNA is 12.1 ± 0.4 ms (Fig. S4). The residential time of Cas9 on each non-PAM binding sites is subsequently estimated as 0.4 ± 0.1 ms. A recent study indicated that a single Cas9 protein spends 6 hours to find a single target site in *E. Coli*, whose genome size is about 5×10^6^ bp (Jones et al., 2017). Based on our measured value, a single Cas9 only takes about 0.5 hours to one-dimensionally diffuse through PAM-free dsDNA of the same size. The discrepancy between these two values emphasizes the complexity of target search of Cas9 in cell. Extra time is spent by Cas9 on interactions with enormous numbers of PAM sites and off-target sites, crowded cellular environment and with random searching processes.

Although 1D diffusion of Cas9 does not require external energy, we captured a surprisingly biased 1D diffusion of Cas9 on dsDNA under physiological buffer condition. Low-salt concentration, which strengthens interactions between DNA binding proteins and dsDNA (Berg et al., 1981; Bonnet et al., 2008; Lohman, 1986; Mondal and Bhattacherjee, 2015; Tempestini et al., 2018), causes Cas9 to display unbiased 1D diffusion. In addition, Cas9-VQR with altered PAM preference exhibits biased diffusion while Cas9-EQR unbiased diffuses on dsDNA under physiological salt condition (Fig. S3, Table S5). The D1135V and D1135E mutations, which are the only different sites between Cas9-VQR and Cas9-EQR, are proposed to play auxiliary roles to accommodate the PAM duplex in a displaced conformation (Anders et al., 2016; Hirano et al., 2016). The fourth G of the PAM sequence in the Cas9-EQR is fixed more tightly than that in the Cas9-VQR, because the aliphatic portion of the D1135E side chain forms a van der Waals contact with its ribose moiety. Together, we proposed that, loose interactions between Cas9 and dsDNA is likely to promote biased 1D diffusion, whereas tight Cas9-dsDNA interactions caused by low-salt concentration or mutations on PAM recognition residues alter energy landscape of diffusion to induce unbiased diffusion.

In summary, we provided a quantitative kinetic scheme to describe how Cas9 combines 3D and biased 1D diffusion to search for its target with high efficiency. Our results emphasized the importance of ionic strength and accessibility of tens of base pairs adjacent to target sites, which dominate target search mechanism and efficiency, respectively. Our findings provided important guidance for both in-vivo and in-vitro applications of Cas9.

## Methods and Materials

### Cas9 purification and labeling

*S. pyogenes* Cas9 (SpCas9) were expressed in *Escherichia coli* strain BL21 (DE3) using the expression plasmid pMJ806 (Addgene plasmid # 39312) (Jinek et al., 2012) and purified as previously described (Yang et al., 2018). Briefly, the cells were sonicated on ice and further clarified by centrifugation. The supernatant was first bound to Ni-NTA agarose (Qiagen) in lysis buffer (20 mM HEPES pH 7.5, 500 mM NaCl, 1 mM TCEP, 1 mM PMSF, 10% glycerol), and then eluted with 20 mM HEPES pH 7.5, 250 mM KCl, 1 mM TCEP, and 150 mM imidazole. After removing the His_6_-MBP affinity tag by overnight digestion at 4°C with TEV protease, the protein was further purified by cation exchange with Source S column (GE Healthcare), eluted with a linear KCl gradient of 0.1 – 1 M KCl, followed by gel filtration chromatography (Superdex 200 Increase 5/150 GL, GE Heathcare) with 50 mM Tris-HCl pH 7.5, 150 mM KCl, 1 mM TCEP, and finally concentrated to 5∼10 mg/mL. The SpCas9 variants designed for site-specific labeling were constructed by QuikChange site-directed mutagenesis with Q5® High-Fidelity DNA Polymerase (NEB) and confirmed by DNA sequencing. The variants were purified as described for the wild type proteins.

The SpCas9 variants were labeled as previously described (Yang et al., 2018). The labeling reactions were conducted in 50 mM Tris-HCl pH 7.5, 150 mM KCl, 1 mM TCEP containing 30 mM Cas9, 300 mM maleimide-Cy3 and 500 mM malemide-Cy5 (LumiProbe). After 2 hours incubation at room temperature (∼25 °C), reactions were quenched by adding 10 mM DTT, and excess free dyes were removed by Sephadex G-25 column (Nap-5, GE Healthcare). Labeling efficiency was estimated by measuring A_280_, A_549_, and A_649_. Proteins were all flash-frozen in liquid nitrogen and stored at −80°C.

### *In-vitro* transcription and purification of RNA

The sgRNA was transcribed and purified as previously described (Yang et al., 2018). DNA templates carrying T7 promoter and sgRNA sequence were cloned into a pMV Vector. The plasmid was linearized with BamHI digestion and further recovered as a template for in vitro transcription via HiScribe™ T7 High Yield RNA Synthesis Kit (NEB). RNA was isolated by 15% denaturing PAGE and dissolved in stocking buffer (300 mM NaAc pH 5.2, 1 mM EDTA) at 55°C. The soluble fraction containing RNA is recovered by ethanol precipitation. The integrity and purity of RNA were confirmed by 1.2% agarose gel electrophoresis. The purified RNA sample was dissolved in nuclease-free H_2_O (Invitrogen).

### *In-vitro* cleavage assay

The target-containing plasmid (Table S1) used in Fig. 1a was purchased from Qinglan Biotech (Jiangsu, China) and linearized into two fragments by PCR amplification, purified and recovered from 0.8∼2% agarose gel. Primers (Table S1) used for linearization were commercially synthesized by Sangon Biotech (Shanghai, China). DNA substrates were prepared by mixing equimolar amounts of two linearized fragments in reaction buffer (50 mM Tris-HCl pH 7.5, 150 mM KCl, 5 mM MgCl_2_, 1 mM DTT). sgRNA was pre-heated in reaction buffer at 95°C for 5 min and slowly cooling down to room temperature. SpCas9/sgRNA complexes were reconstituted by mixing SpCas9 with a 4× molar excess of the sgRNA in reaction buffer and incubating at 25°C for 20 minutes. Cleavage assay were initiated at the time point when adding Cas9/sgRNA complex (final concentration 28 nM) into two linearized DNA fragments (final concentration 7 nM), and terminated by adding 6× DNA loading buffer (NEB). For cleavage assay presented in Figs. 1d, e, and f, the reaction containing DNA substrates with a final concentration of 10 nM (molar ratio of DNA:Cas9/sgRNA=1:2). Percentage of cleaved DNA was resolved by agarose electrophoresis and apparent cleavage rates were extracted by single exponential curves. S.E.M. were estimated through three or more replicates.

Primers (Table S1) for short target-containing fragments amplification (≤240 bp) were purchased from Sangon Biotech (Shanghai, China) with 5’ amino modification, which were labeled by mixing N-hydroxysuccinimido (NHS) Cy5 and DNAs at 20:1 molar ratio at 37 °C for 4 h. Labeled DNAs were separated from excess free fluorophores through ethanol precipitation. Then, the short target-containing fragments were obtained by PCR amplification or directed annealing with labeled primers. The cleavage reactions were performed with 7 nM DNAs (molar ratio of DNA:Cas9/sgRNA=1:4) in cleavage buffer and quenched by 2 × formamide loading dye and resolved on 12.5% denaturing polyacrylamide gels containing 8 M urea and visualized by GE typhoon FLA9500 imaging system.

### Acquisition of single-molecule fluorescent data

All smFRET experiments were performed at 25 °C in the reaction imaging buffer (50 mM Tris-HCl pH 7.5, 100 mM KCl, 5 mM MgCl_2_, 1 mM DTT, 3 mg/mL glucose, 100 µg/mL glucose oxidase (Sigma-Aldrich), 40 μg/mL catalase (Roche), 1 mM cyclooctatetraene (COT, Sigma-Aldrich), 1 mM 4-nitrobenzylalcohol (NBA, Sigma-Aldrich), 1.5 mM 6-hydroxy-2,5,7,8-tetramethyl-chromane-2-carboxylic acid (Trolox, Sigma-Aldrich – added from a concentrated DMSO stock solution)). Single-molecule spectroscopic microscopy was performed on a home-built objective-type TIRF microscope, based on a Nikon Eclipse Ti-E with an EMCCD camera (Andor iXon Ultra 897), and solid state 532 nm excitation lasers (Coherent Inc. OBIS Smart Lasers) (Peng et al., 2017). Fluorescence emission from the probes was collected by the microscope and spectrally separated by interference dichroic (T635lpxr, Chroma) and bandpass filters, ET585/65m (Chroma, Cy3) and ET700/75m (Chroma, Cy5), in a Dual-View spectral splitter (Photometrics, Inc., Tucson, AZ). Hardwares were controlled and single molecule movies were collected using Cell Vision software (Beijing Coolight Technology).

### Single-molecule fluorescent experiments

Biotinylated DNA strands and non-modified DNA strands used for single-molecule fluorescent assay (Table S3) were purchased from Sangon Biotech (Shanghai, China). DNA substrates were prepared by mixing equimolar amounts of biotinylated and non-modified DNA strands (≤ 60 bp) heating at 65°C for 1-2 min and then slowly cooling down to room temperature, or by PCR amplification (> 60 bp). 200 pM biotinylated DNA was firstly immobilized to streptavidin-decorated slides. The excessive unbound DNAs were removed by washing with reaction imaging buffer. Labeled Cas9/sgRNA (100 nM: 400 nM) were pre-incubated at 25 °C in reaction buffer (50 mM Tris-HCl pH 7.5, 150 mM KCl, 5 mM MgCl_2_, 1 mM DTT). To record the appearance time of Cas9/sgRNA, the collection of movies began 5 s prior to injection of 50 µL pre-incubated 5 nM labeled Cas9/sgRNA complex in reaction imaging buffer and carried out without further washing. The association rates of Cas9/sgRNA towards different DNA substrates were determined by recording each appearance time upon individual stable binding event occurred and fitted with single exponential decay.

For single-molecule fluorescent experiments at low salt conditions, the reaction was carry out in low-salt reaction buffer (50 mM HEPES-NaOH pH 7.5, 10 mM NaCl, 2 mM MgCl_2_, 1 mM DTT)

## Acknowledgement

This project was supported by funds from the National Natural Science Foundation of China (21877069, 21922704 and 31570754), Tsinghua-Peking Joint Center for Life Sciences, Beijing Advanced Innovation Center for Structural Biology and Beijing Frontier Research Center for Biological Structure to CC.

## Author Contributions

MY and CC designed the experiments; MY, LZ, RS, WZ and YY prepared materials and reagents and performed experiments; MY and CC analyzed the data and wrote the paper.

