## Supplemental Figures and Tables for "Streptococcus pyogenes Cas9 displays biased one-dimensional diffusion on dsDNA to search for a target"

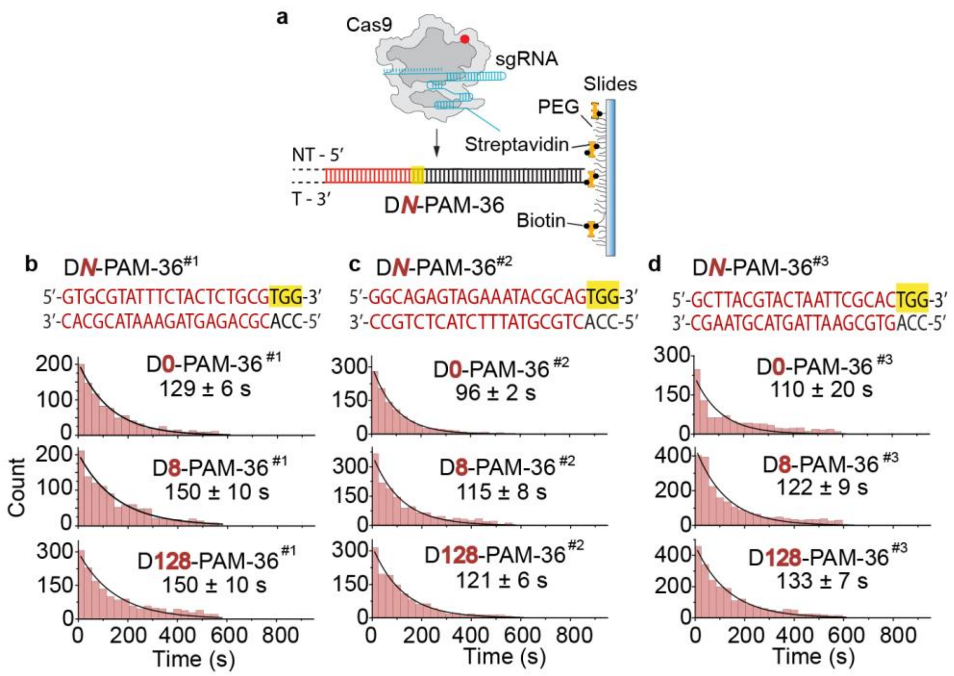


**Figure S1. Single-molecule fluorescence assays on three different DNA series containing distinct target sites. Related to Figure 3.**

**a)** Scheme of single-molecule assay to capture apparent target search rates of Cas9 on D*N*-PAM-36 DNAs. **b-d)** Distributions of appearance time of Cas9 on three series of D*N*-PAM-36 DNAs. D*N*-PAM-36$^{\#1}$ (**b**), D*N*-PAM-36$^{\#2}$ (**c**), and D*N*-PAM-36$^{\#3}$ (**d**) containing different target sites, which were listed and colored in red. Single exponential decay was used to extract average appearance time from injection until stable binding of Cas9 on dsDNAs.

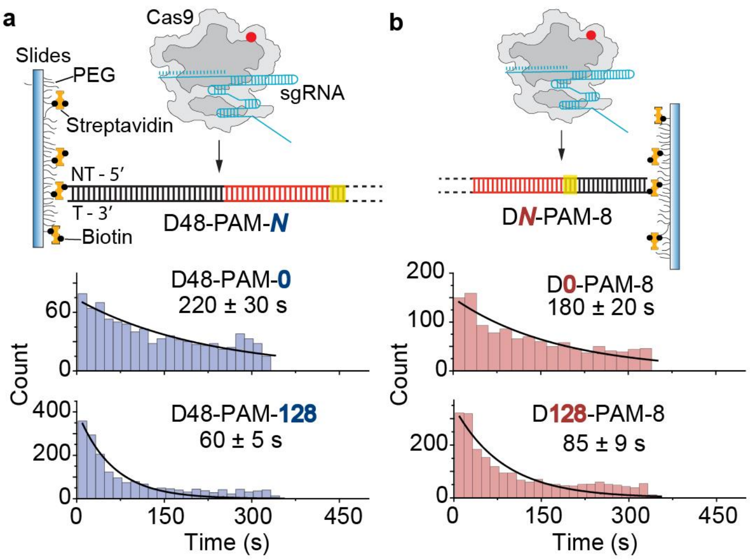


**Figure S2. Unbiased 1D diffusion under low salt condition. Related to Figure 3.**

a) Scheme of single-molecule assay and distributions of appearance time of Cas9 on D48-PAM-*N* DNAs in low salt buffer (50 mM HEPES-NaOH pH 7.5, 10 mM NaCl, 2 mM $MgCl_2$, 1 mM DTT). b) Scheme of single-molecule assay and distributions of appearance time of Cas9 on D*N*-PAM-8 DNAs in low satl buffer. Single exponential decay was used to extract average appearance time from injection until stable binding of Cas9 on dsDNAs.

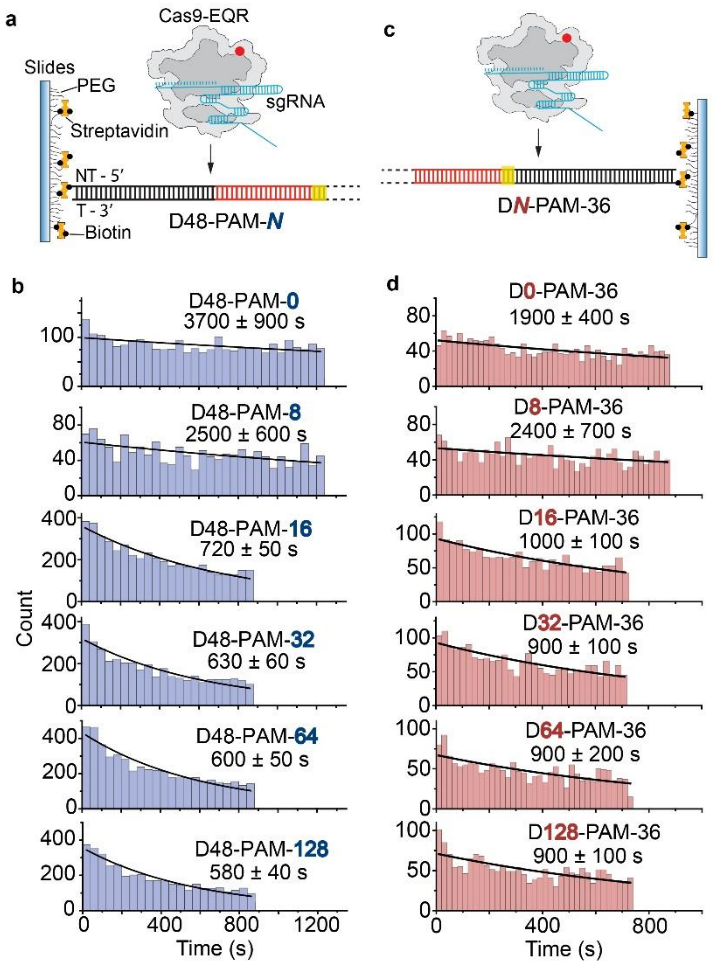


**Figure S3 Single molecule fluorescence assay of Cas9-EQR. Related to Figure 3.**

**a)** Scheme of single-molecule fluorescence assay to capture apparent target search rates of Cas9-EQR on D48-PAM-*N* DNAs. The PAM sequence is 5’-TGAG-3’ that is recognized by Cas9-EQR. **b)** Distributions of appearance time of Cas9-EQR on D48-PAM-*N* DNAs, which were fitted by single exponential decay. **c)** Scheme of capturing apparent target search rates of Cas9-EQR on D*N*-PAM-36 DNAs. **d)** Distributions of appearance time of Cas9-EQR on D*N*-PAM-36 DNAs.

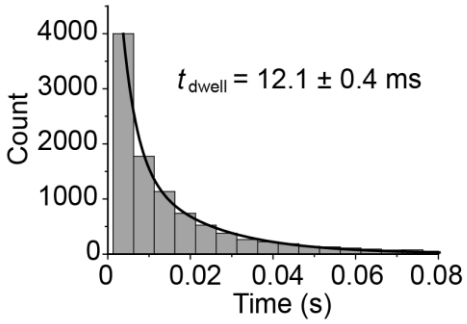


**Figure S4. Dwell time distributions of Cas9/sgRNA on a 59-bp dsDNA containing no PAM and no target sites. Related to Figure 3.** 2.5 ms per frame was used to capture transient interactions. The weighted average residence time was extracted using double exponential decay.

**Table S1. Plasmid and primer sequences for *in-vitro* cleavage assay. Related to Figures 1 and 2.**

| **Name** [a] | **Sequences** [b] |
|---|---|
| Target plasmid | AAAAATAAACAAATAGGGGTTCCGCGCACATTTCCCCGAAAAGTGCCA<br>CCTGACGTCTAAGAAACCATTATTATCATGACATTAACCTATAAAAATA<br>GGCGTATCACGAGGCCCTTTCGTTGTAAAACGACGGCCAGTCCGTCTCT<br>ATCCGGTCTCGATCCGCAGTCTCTTGGCACAGGTCTAGAGACGAGCAG<br>AAATCTCTGCTGACGCATAAAGATGAGACGCTGGAGTACAAACGTCAG<br>CTCATATGACCGTGCGAGTTACTGCCAACCGAGACCCAACCGAGACGG<br>GTCATAGCTGTTTCCAGTGTGCCGCTTCCTCGCTCACTGACTCGCTGCG<br>CTCGGTCGTTCGGCTGCGGCGAGCGGTATCAGCTCACTCAAAGGCGGT<br>AATACGGTTACCCACAGAATCAGGGGATAACGCAGGAAAGAACATGT<br>GAGCAAAAGGCCAGCAAAAGGCCAGGAACCGTAAAAAGGCCGCGTTG<br>CTGGCGTTTTTCCATAGGCTCCGCCCCCCTGACGAGCATCACAAAAATC<br>GACGCTCAAGTCAGAGGTGGCGAAACCCGACAGGACTATAAAGATACC<br>AGGCGTTTCCCCCTGGAAGCTCCCTCGTGCGCTCTCCTGTTCCGACCCT<br>GCCGCTTACCGGATACCTGTCCGCCTTTCTCCCTTCGGGAAGCGTGGCG<br>CTTTCTCAATGCTCACGCTGTAGGTATCTCAGTTCGGTGTAGGTCGTTC<br>GCTCCAAGCTGGGCTGTGTGCACGAACCCCCCGTTCAGCCCGACCGCT<br>GCGCCTTATCCGGTAACTATCGTCTTGAGTCCAACCCGGTAAGACACGA<br>CTTATCGCCACTGGCAGCAGCCACTGGTAACAGGATTAGCAGAGCGAG<br>GTATGTAGGCGGTGCTACAGAGTTCTTGAAGTGGTGGCCTAACTACGG<br>CTACACTAGAAGGACAGTATTTGGTATCTGCGCTCTGCTGAAGCCAGTT<br>ACCTTCGGAAAAAGAGTTGGTAGCTCTTGATCCGGCAAACAAACCACC<br>GCTGGTAGCGGTGGTTTTTTTGTTTGCAAGCAGCAGATTACGCGCAGAA<br>AAAAAGGATCTCAAGAAGATCCTTTGATCTTTTCTACGGGGTCTGACGC<br>TCAGTGGAACGAAAACTCACGTTAAGGGATTTTGGTCATGAGATTATC<br>AAAAAGGATCTTCACCTAGATCCTTTTAAATTAAAAATGAAGTTTTAAA<br>TCAATCTAAAGTATATATGAGTAAACTTGGTCTGACAGTTACCAATGCT<br>TAATCAGTGAGGCACCTATCTCAGCGATCTGTCTATTTCGTTCATCCAT<br>AGTTGCCTGACTCCCCGTCGTGTAGATAACTACGATACGGGAGGGCTT<br>ACCATCTGGCCCCAGTGCTGCAATAATACCGCGGGACCCACGCTCACC<br>GGCTCCAGATTTATCAGCAATAAACCAGCCAGCCGGAAGGGCCGAGCG<br>CAGAAGTGGTCCTGCAACTTTATCCGCCTCCATCCAGTCTATTAATTGT<br>TGCCGGGAAGCTAGAGTAAGTAGTTCGCCAGTTAATAGTTTGCGCAAC<br>GTTGTTGCCATCGCTACAGGCATCGTGGTGTCACGCTCGTCGTTTGGTA<br>TGGCTTCATTCAGCTCCGGTTCCCAACGATCAAGGCGAGTTACATGATC<br>CCCCATGTTGTGCAAAAAAGCGGTTAGCTCCTTCGGTCCTCCGATCGTT<br>GTCAGAAGTAAGTTGGCCGCCGTGTTATCACTCATGGTTATGGCAGCAC<br>TACATAATTCTCTTACTGTCATGCCATCCGTAAGATGCTTTTCTGTGACT<br>GGTGAGTACTCAACCAAGTCATTCTGAGAATAGTGTATGCGGCGACCG<br>AGTTGCTCTTGCCCGGCGTCAATACGGGATAATACCGCGCCACATAGC<br>AGAACTTTAAAAGTGCTCATCATTGGAAAACGTTCTTCGGGGCGAAAA<br>CTCTCAAGGATCTTACCGCTGTTGAGATCCAGTTCGATGTAACCCACTC<br>GTGCACCCAACTGATCTTCAGCATCTTTTACTTTCACCAGCGTTTCTGG<br>GTGAGCAAAAACAGGAAGGCAAAATGCCGCAAAAAAGGGAATAAGGG<br>CGACACGGAAATGTTGAATACTCATACTCTTCCTTTTTCAATATTATTG<br>AAGCATTTATCAGGGTTATTGTCTCATGAGCGGATACATATTTGAATGT<br>ATTTAG |
| 2188bp-F | TAGTTGCCTGACTCCCCGTCGT |
| 2188bp-R | TGGATGAACGAAATAGACAGATCGCTGAG |
| 30bp-F | GACGCATAAAGATGAGACGCTGGAGTACAAA |
| 30bp-R | TTTGTACTCCAGCGTCTCATCTTTATGCGTC |
| Vector-30-F | CGTCAGCTCATATGACCGTGC |
| Vector-30-R | AGCAGAGATTTCTGCTCGTCTCTAG |

| | |
|---|---|
| 59bp-F | GACGAGCAGAAATCTCTGCTGACGCATAAAGATGAGACGCTGGAGTAC<br>AAACGTCAGCT |
| 59bp-R | AGCTGACGTTTGTACTCCAGCGTCTCATCTTTATGCGTCAGCAGAGATT<br>TCTGCTCGTC |
| Vector-59-F | CATATGACCGTGCGAGTTACTGCCAA |
| Vector-59-R | TCTAGACCTGTGCCAAGAGACTGC |
| 120bp-F | GATCCGCAGTCTCTTGGCAC |
| 120bp-R | GTCTCGGTTGGCAGTAACTCG |
| Vector-120-F | CCAACCGAGACGGGTCATAGC |
| Vector-120-R | GAGACCGGATAGAGACGGACTGG |
| 240bp-F | GGCGTATCACGAGGCCCTTTC |
| 240bp-R | CAGCGAGTCAGTGAGCGAGG |
| vector-240-F | CGCTCGGTCGTTCGGCTGCG |
| vector-240-R | ATAGGTTAATGTCATGATAATAATGGTTTCTTAGACGT |
| 480bp-F | GGATACATATTTGAATGTATTTAGAAAAATAAAC |
| 480bp-R | GGCCTTTTGCTGGCCTTTTGC |
| vector-480-F | AGGAACCGTAAAAAGGCCGCG |
| vector-480-R | GCTCATGAGACAATAACCCTGATAAATGCTTC |
| 960bp-F | CTTCGGGGCGAAAACTCTCAAG |
| 960bp-R | CAGCGTGAGCATTGAGAAAGCG |
| vector-960-F | TAGGTATCTCAGTTCGGTGTAGGTCGTTCG |
| vector-960-R | AACGTTTTCCAATGATGAGCACTTTTAAAGTTC |
| Substrate #1 | AGACGTGACGAGTTACTGTGTATCAAGCGAGAGTCAAGCGAGACGTCG<br>TCATAGCTGTTTGCAGTGTGCGCGCTTGCTCGCTCACTGACTCGCTGCG<br>CTCGCTCGTTCGTGCTGCGCGAGCGCTATCAGCTCACTCAAAGCGCTAA<br>TACGCTTAGCACAGAATCAGCGCATAACGCAGCAAAGAACATGTGAGC<br>AAAAGCTAGCAAAAGCGAGCAAGCGTAAAAAGCGCGTTGCTCGCGTAT<br>AGTAATGAACAGCAATGTTTAGACTTAAGCGATGCTTGCATCTCATAAT<br>CTGACAGACGCATAAAGATGAGACGCTGGTCGATCATTTTACTGACTA<br>TGAGTGCAAAACGAATCACGATAGCTAAAGATAATTGATTTTGCATAC<br>GCTGCTGCGCGCTGACGAGCATCACAAAAATCGACGCTCAAGTCAGAG<br>CTGCGAAAGCTGACAGCACTATAAAGATAGCAGCGTTTGCGCTGCAAG<br>CTGCTCGTGCGCTCTGCTGTTGCGAGCTGCGCTTAGCGCATAGCTGTGC<br>TGCTTTCTGCTTCGCGAAGCGTGCGCTTTCTCAATGCTCACGCTGTAGC<br>TATCTCAGTTCGCTGTAAACTGTCTGACAGTACGAT |
| Substrate #2 | AGACGTGACGAGTTACTGTGTATCAAGCGAGAGTCAAGCGAGACGTCG<br>TCATAGCTGTTTGCAGTGTGCGCGCTTGCTCGCTCACTGACTCGCTGCG<br>CTCGCTCGTTCGTGCTGCGCGAGCGCTATCAGCTCACTCAAAGCGCTAA<br>TACGCTTAGCACAGAATCAGCGCATAACGCAGCAAAGAACATGTGAGC<br>AAAAGCTAGCAAAAGCGAGCAAGCGTAAAAAGCGCGTTGCTCGCGTAT<br>AGTAATGGACAGCAATGGTTAGACTTGGGCGATGCTGGCATCTCATGG<br>TCTGACAGACGCATAAAGATGAGACGCTGGTCGATCATGGTACTGACT<br>GGGAGTGCATGGCGAATCATGGTAGCTAATGGTAATTGATTTTGCATA<br>CGCTGCTGCGCGCTGACGAGCATCACAAAAATCGACGCTCAAGTCAGA<br>GCTGCGAAAGCTGACAGCACTATAAAGATAGCAGCGTTTGCGCTGCAA<br>GCTGCTCGTGCGCTCTGCTGTTGCGAGCTGCGCTTAGCGCATAGCTGTG<br>CTGCTTTCTGCTTCGCGAAGCGTGCGCTTTCTCAATGCTCACGCTGTAG<br>CTATCTCAGTTCGCTGTAAACTGTCTGACAGTACGAT |

| | |
|---|---|
| Substrate #3 | AGACGTGACGAGTTACTGTGTATCAAGCGAGAGTCAAGCGAGACGTCG<br>TCATAGCTGTTTGCAGTGTGCGCGCTTGCTCGCTCACTGACTCGCTGCG<br>CTCGCTCGTTCGTGCTGCGCGAGCGCTATCAGCTCACTCAAAGCGCTAA<br>TACGCTTAGCACAGAATCAGCGCATAACGCAGCAAAGAACATGTGAGC<br>AAAAGCTAGCAAAAGCGAGCAAGCGTAAAAAGCGCGTTGCTCGCGT<br>TGTCAGACCATGAGATGCCAGCATCGCCCAAGTCTAACCATTGCTGT<br>CCATTACTATGACGCATAAAGATGAGACGCTGGTCAATTACCATTAG<br>CTACCATGATTCGCCATGCACTCCCAGTCAGTACCATGATCGA<br>TTTTGCATACGCTGCTGCGCGCTGACGAGCATCACAAAAATCGACGCTC<br>AAGTCAGAGCTGCGAAAGCTGACAGCACTATAAAGATAGCAGCGTTTG<br>CGCTGCAAGCTGCTCGTGCGCTCTGCTGTTGCGAGCTGCGCTTAGCGCA<br>TAGCTGTGCTGCTTTCTGCTTCGCGAAGCGTGCGCTTTCTCAATGCTCA<br>CGCTGTAGCTATCTCAGTTCGCTGTAAACTGTCTGACAGTACGAT |
| D8-PAM-28 | ATCTGACAGACGCATAAAGATGAGACGCTGGTCGATCATTTTACTGAC<br>TATGAGTGCAA |
| D20-PAM-16 | GACGAGCAGAAATCTCTGCTGACGCATAAAGATGAGACGCTGGAGTAC<br>AAACGTCAGCT |
| D28-PAM-8 | AGCGATGCTTGCATCTCATAATCTGACAGACGCATAAAGATGAGACGC<br>TGGTCGATCAT |
| D0-PAM-36[as] | GGCAGAGTAGAAATACGCAGTGGAGTACAAACGTCAGCTCATATGACCG<br>TGCGAGTTAC |
| D20-PAM-16[as] | GACGAGCAGAAATCTCTGCTGGCAGAGTAGAAATACGCAGTGGAGTAC<br>AAACGTCAGCT |
| D28-PAM-8[as] | GGTCTAGAGACGAGCAGAAATCTCTGCTGGCAGAGTAGAAATACGCAG<br>TGGAGTACAAA |

[a] as stands for alternative sequence.
[b] Sequences are listed from 5’-3’. Only non-target strands of dsDNA are shown. Protospacer sequence is colored in red and PAM sequence is 5’-TGG-3’ and highlighted in yellow. Underline sequences indicate the position containing PAM at the complementary target strands.

**Table S2. Cleavage rates of *in-vitro* cleavage assay. Related to Figures 1 and 2.**

| Target dsDNA groups | Cleavage rate($10^{-3}$ $s^{-1}$) |
|---|---|
| D30 | 2.8 ± 0.5 |
| D59 | 6.2 ± 0.7 |
| D120 | 8.3 ± 0.5 |
| D240 | 8.3 ± 0.5 |
| D480 | 9.8 ± 0.5 |
| D960 | 9.3 ± 0.5 |
| D2188 | 11.2 ± 0.7 |
| Substrate #1 | 8.2 ± 0.8 |
| Substrate #2 | 6.2 ± 0.8 |
| Substrate #3 | 3.7 ± 0.5 |
| D8-PAM-28 | 17 ± 2 |
| D20-PAM-16 | 7.2 ± 0.2 |
| D28-PAM-8 | 3.1 ± 0.2 |
| D0-PAM-36$^{as}$ | 22 ± 2 |
| D20-PAM-16$^{as}$ | 8.3 ± 1.3 |
| D28-PAM-8$^{as}$ | 4.7 ± 0.8 |

All results were averages of three repeated experiments.

**Table S3. Sequence of sgRNA and DNA substrates used for single-molecule fluorescence assay. Related to Figures 3 and S1-S4.**

| Description | Sequences [a] |
|---|---|
| sgRNA | 5’-GACGCAUAAAGAUGAGACGCGUUUUAGAGCUAUGCUGUUUUGGAAACAAAACAGC AUAGCAAGUUAAAAUAAGGCUAGUCCGUUAUCAACUUGAAAAAGUGGCACCGAGUG GUGCUUUUUUU-3’ |
| sgRNA#1 | 5’-GUGCGUAUUUCUACUCUGCGGUUUUAGAGCUAUGCUGUUUUGGAAACAAAACAGC AUAGCAAGUUAAAAUAAGGCUAGUCCGUUAUCAACUUGAAAAAGUGGCACCGAGUG GUGCUUUUUUU-3’ |
| sgRNA#2 | 5’-GGCAGAGUAGAAAUACGCAGGUUUUAGAGCUAUGCUGUUUUGGAAACAAAACAG CAUAGCAAGUUAAAAUAAGGCUAGUCCGUUAUCAACUUGAAAAAGUGGCACCGAGU GGUGCUUUUUUU-3’ |
| sgRNA#3 | 5’-GCUUACGUACUAAUUCGCACGUUUUAGAGCUAUGCUGUUUUGGAAACAAAACAGC AUAGCAAGUUAAAAUAAGGCUAGUCCGUUAUCAACUUGAAAAAGUGGCACCGAGUG GUGCUUUUUUU-3’ |
| D48-PAM-0 | 5’-Biotin-AGCGATGCTTGCATCTCATAATCTGACAGACGCATAAAGATGAGACGCTGG-3’ |
| D48-PAM-8 | 5’-Biotin-AGCGATGCTTGCATCTCATAATCTGACAGACGCATAAAGATGAGACGCTGGTC GATCAT-3’ |
| D48-PAM-16 | 5’-Biotin-AGCGATGCTTGCATCTCATAATCTGACAGACGCATAAAGATGAGACGCTGGTC GATCATTTTACTGA-3’ |
| D48-PAM-32 | 5’-Biotin-AGCGATGCTTGCATCTCATAATCTGACAGACGCATAAAGATGAGACGCTGGTC GATCATTTTACTGACTATGAGTGCAAAACG-3’ |
| D48-PAM-64 | 5’-Biotin-AGCGATGCTTGCATCTCATAATCTGACAGACGCATAAAGATGAGACGCTGGTC GATCATTTTACTGACTATGAGTGCAAAACGAATCACGATAGCTAAAGATAATT GATTTTGCA-3’ |
| D48-PAM-128 | 5’-Biotin-AGCGATGCTTGCATCTCATAATCTGACAGACGCATAAAGATGAGACGCTGGTC GATCATTTTACTGACTATGAGTGCAAAACGAATCACGATAGCTAAAGATAATT GATTTTGCATACGCTGCTGCGCGCTGACGAGCATCACAAAAATCGACGCTCAA GTCAGAGCTGCGAAAGCTGA-3’ |
| D0-PAM-36 | 5’-GACGCATAAAGATGAGACGCTGGTCGATCATTTTACTGACTATGAGTGCAAAACGAA TC-Biotin-3’ |
| D8-PAM-36 | 5’-ATCTGACAGACGCATAAAGATGAGACGCTGGTCGATCATTTTACTGACTATGAGTGC AAAACGAATC-Biotin-3’ |
| D16-PAM-36 | 5’-ATCTCATAATCTGACAGACGCATAAAGATGAGACGCTGGTCGATCATTTTACTGACTA TGAGTGCAAAACGAATC-Biotin-3’ |
| D32-PAM-36 | 5’-CTTAAGCGATGCTTGCATCTCATAATCTGACAGACGCATAAAGATGAGACGCTGGTC GATCATTTTACTGACTATGAGTGCAAAACGAATC-Biotin-3’ |
| D64-PAM-36 | 5’-CTCGCGTATAGTAATGAACAGCAATGTTTAGACTTAAGCGATGCTTGCATCTCATAAT CTGACAGACGCATAAAGATGAGACGCTGGTCGATCATTTTACTGACTATGAGTGCAAAA CGAATC-Biotin-3’ |
| D128-PAM-36 | 5’-ATAACGCAGCAAAGAACATGTGAGCAAAAGCTAGCAAAAGCGAGCAAGCGTAAAAA GCGCGTTGCTCGCGTATAGTAATGAACAGCAATGTTTAGACTTAAGCGATGCTTGCATCT CATAATCTGACAGACGCATAAAGATGAGACGCTGGTCGATCATTTTACTGACTATGAGT GCAAAACGAATC-Biotin-3’ |
| D0-PAM-8 | 5’-GACGCATAAAGATGAGACGCTGGTCGATCAT-Biotin-3’ |
| D8-PAM-8 | 5’-ATCTGACAGACGCATAAAGATGAGACGCTGGTCGATCAT-Biotin-3’ |
| D128-PAM-8 | 5’-ATAACGCAGCAAAGAACATGTGAGCAAAAGCTAGCAAAAGCGAGCAAGCGTAAAAA GCGCGTTGCTCGCGTATAGTAATGAACAGCAATGTTTAGACTTAAGCGATGCTTGCAT CTCATAATCTGACAGACGCATAAAGATGAGACGCTGGTCGATCAT-Biotin-3’ |
| No-PAM-No-Target | 5’-GACGAGCTGCATAACGCGAAAAAATATATTTATCTGCTTGATCTTCAAATGTTGTATTG-3 |

[a] sgRNA guide sequence and target sites are shown in red. #1, #2 and #3 refer to 3 different spacer sequences. PAM sequence is 5’-TGG-3’ and highlighted in yellow. Biotinylated DNA duplexes are named referring to the target and PAM location relative to the 5’ and 3’ end of the NT strand, and only non-target strand sequence is listed in this table. DNAs containing full-cognate target sequences paired with the spacers in sgRNA#1, sgRNA#2 or sgRNA#3 are designed based on the same principles and are not listed for clarity.

**Table S4. Apparent target search rates of Cas9/sgRNA under different salt concentrations. Related to Figures 3, S1 and S2.**

| Buffer | DNA substrates | Appearance rates of Cas9/sgRNA ($\mu M^{-1}$ $s^{-1}$) |
|---|---|---|
| Buffer 1 | D48-PAM-0 | 0.20 ± 0.02 |
| | D48-PAM-8 | 0.36 ± 0.02 |
| | D48-PAM-16 | 0.88 ± 0.04 |
| | D48-PAM-32 | 1.36 ± 0.04 |
| | D48-PAM-64 | 1.84 ± 0.08 |
| | D48-PAM-128 | 1.72 ± 0.08 |
| | D0-PAM-36 | 0.96 ± 0.08 |
| | D8-PAM-36 | 0.84 ± 0.06 |
| | D16-PAM-36 | 0.68 ± 0.06 |
| | D32-PAM-36 | 0.74 ± 0.04 |
| | D64-PAM-36 | 0.9 ± 0.1 |
| | D128-PAM-36 | 0.72 ± 0.06 |
| | D0-PAM-8 | N.A |
| | D8-PAM-8 | N.A |
| | D128-PAM-8 | N.A |
| | D0-PAM-36#1 | 1.54 ± 0.06 |
| | D8-PAM-36#1 | 1.3 ± 0.1 |
| | D128-PAM-36#1 | 1.4 ± 0.1 |
| | D0-PAM-36#2 | 2.08 ± 0.04 |
| | D8-PAM-36#2 | 1.74 ± 0.12 |
| | D128-PAM-36#2 | 1.65 ± 0.08 |
| | D0-PAM-36#3 | 1.92 ± 0.28 |
| | D8-PAM-36#3 | 1.64 ± 0.12 |
| | D128-PAM-36#3 | 1.52 ± 0.08 |
| Buffer 2 (low salt) | D0-PAM-8 | 1.14 ± 0.16 |
| | D128-PAM-8 | 2.4 ± 0.2 |
| | D48-PAM-0 | 0.9 ± 0.1 |
| | D48-PAM-128 | 3.3 ± 0.3 |

Buffer 1: 50 mM Tris-HCl pH 7.5, 150 mM KCl, 5 mM $MgCl_2$, 1 mM DTT; Buffer 2: 50 mM HEPES-NaOH pH 7.5, 10 mM NaCl, 2 mM $MgCl_2$, 1 mM DTT. All results were averages of three repeated experiments. #1, #2 and #3 refer to 3 different spacer sequences.

**Table S5. Apparent target search rates of Cas9 variants which recognized alternative PAM sequences. Related to Figures 3 and S3.**

| Cas9 variants | DNA substrates | Appearance rates of Cas9/sgRNA ($\mu M^{-1}$ $s^{-1}$) |
|---|---|---|
| Cas9-EQR | D48-PAM-0 | 0.0027 ± 0.0006 |
| | D48-PAM-8 | 0.004 ± 0.001 |
| | D48-PAM-16 | 0.014 ± 0.001 |
| | D48-PAM-32 | 0.016 ± 0.002 |
| | D48-PAM-64 | 0.017 ± 0.001 |
| | D48-PAM-128 | 0.017 ± 0.001 |
| | D0-PAM-36 | 0.005 ± 0.01 |
| | D8-PAM-36 | 0.004 ± 0.01 |
| | D16-PAM-36 | 0.011 ± 0.001 |
| | D32-PAM-36 | 0.013 ± 0.001 |
| | D64-PAM-36 | 0.013 ± 0.01 |
| | D128-PAM-36 | 0.013 ± 0.001 |
| Cas9-VQR | D48-PAM-0 | 0.05 ± 0.01 |
| | D48-PAM-8 | 0.09 ± 0.01 |
| | D48-PAM-128 | 0.61 ± 0.01 |
| | D0-PAM-36 | 0.22 ± 0.01 |
| | D128-PAM-36 | 0.22 ± 0.01 |

PAM sequence was designed as 5’-TGAG-3’ for Cas9-EQR and Cas9-VQR variants. All results were averages of three repeated experiments.
